# Effects of an odor background on moth pheromone communication: constituent identity matters more than blend complexity

**DOI:** 10.1101/2020.09.03.281311

**Authors:** Lucie Conchou, Philippe Lucas, Nina Deisig, Elodie Demondion, Michel Renou

## Abstract

Olfaction allows insects to communicate with pheromones even in complex olfactory landscapes. It is generally admitted that, due to the binding selectivity of the receptors, general odorants should weakly interfere with pheromone detection. However, laboratory studies show that volatile plant compounds (VPCs) modulate responses to the pheromone in male moths. We used extracellular electrophysiology and calcium imaging to measure the responses to the pheromone of receptor and central neurons in males *Agrotis ipsilon* while exposed to simple or composite backgrounds of VPCs. Maps of activities were built using calcium-imaging to visualize which areas in antennal lobes (ALs) were affected by VPCs. To mimic a natural olfactory landscape short pheromone puffs were delivered over VPC backgrounds. We chose a panel of VPCs with different chemical structures and physicochemical properties representative of the odorant variety encountered by a moth. We evaluated the intrinsic activity of each VPC and compared the impact of VPC backgrounds at antenna and antennal lobe levels. Then, we prepared binary, ternary and quaternary combinations to determine whether blend activity could be deduced from that of its components. Our data confirm that a VPC background interfere with the moth pheromone system in a dose-dependent manner. Interference with the neuronal coding of pheromone signal starts at the periphery. VPCs showed differences in their capacity to elicit Phe-ORN firing response that cannot be explained by differences in stimulus intensities because we adjusted the source concentrations to vapor pressures. Thus, these differences must be attributed to the selectivity of ORs or any other olfactory proteins. The neuronal network in the ALs, which reformats the ORN-input, did not improve pheromone salience. We postulate that the AL network might have evolved to increase sensitivity and encode for fast changes over a wide range of concentrations, possibly at some cost for selectivity. Comparing three- or four-component blends to binary blends or single compound indicated that a blend showed the activity of its most active compound. Thus, although the diversity of a background might increase the probability of including a VPC interacting with the pheromone system, chemical diversity does not seem to be a prominent factor per se. Global warming is significantly affecting plant metabolism so that the emissions of VPCs and resulting odorscapes are modified. Increase in atmospheric mixing rates of VPCs will change olfactory landscapes which, as confirmed in our study, might impact pheromone communication.

## Introduction

Olfactory communication is essential to insects as it is involved in the identification and the location of vital resources such as a food source, a mate, or an oviposition site. Insects have developed an exquisite olfactory sense in terms of sensitivity, specificity and temporal dynamics. Their olfactory system enables them to discriminate the pheromones they produce, as well as the odors involved in interspecific interactions, such as, for instance, the floral compounds emitted by plants to attract specialist pollinators. Herbivorous species, for instance, can discriminate potential host-plant species based on their volatile emissions (Conchou, Anderson et al. 2017). Exchanges of chemical information are thus not only vital to insects, but also essential to the functioning of the species network composing a community (Kantsa, Raguso et al. 2019). Once released in the atmosphere, these ecologically relevant signals and cues are transported by airflows, diluted, and mixed to a background of other volatile organic compounds to form a complex and changing olfactory landscape (Conchou, Lucas et al. 2019). Considering the hundreds of different volatile compounds released by plants (VPCs) (Knudsen, Tollsten et al. 1993, Knudsen, Eriksson et al. 2006), the capacity of the insect olfactory system to extract the ecologically relevant information from that very complex chemical environment is remarkable.

Insect olfactory systems evolved to deal with such complex olfactory landscapes (Hansson and Stensmyr 2011). Male moths for instance are able to detect from hundreds of meters the odor plume generated by a female emitting its sex pheromone and to navigate upwind toward the calling female. Female moths release a few ng per hour of a specific pheromone blend, which represent only traces compared to the ppb of VPCs present in the atmosphere (Kesselmeier, Kuhn et al. 2000, Wiedenmyer, Friedfeld et al. 2011). In the male antennae, narrowly tuned olfactory receptors (ORs) expressed in olfactory receptor neurons (ORNs) specifically bind the pheromone components (De Bruyne and Baker 2008, Su, Menuz et al. 2009) ensuring detection selectivity. The antennae house thousands of ORNs each of them expressing one functional type of OR specialized in the detection of one pheromone constituent (Phe-ORNs). Beside quality, the firing activities of Phe-ORNs also encode the intensity of the stimulus. Phe-ORNs converge onto a comparatively small number of central neurons in a specialized area of the antennal lobes (AL), the macro-glomerular complex (MGC) (Hansson and Anton 2000). As a consequence of this convergence the projection neurons (PNs) in the MGC display a remarkably low response threshold (Jarriault, Gadenne et al. 2009, Jarriault, Gadenne et al. 2010, Rospars, Gremiaux et al. 2014). Male moths not only discriminate the pheromone constituents, but also show ratio selectivity (Martin, Lei et al. 2013) which increases the specificity of pheromone communication. Blend ratio coding starts in the MGC, some MGC neurons responding more to blend of pheromone constituents than individual components (Anton and Hansson 1995, Wu, Anton et al. 1996). Additionally, fast fluctuations of pheromone concentration are tracked by the ORN and MGC-neurons firing (Gorur-Shandilya, Demir et al. 2017, Jacob, Monsempès et al. 2017, Levakova, Kostal et al. 2018, Levakova, Kostal et al. 2019).

It is generally admitted that, due to the high binding selectivity of the pheromone receptors, general odorants should only weakly interfere with pheromone detection. However, a number of laboratory studies have shown that VPCs modulate pheromone responses in male moths. Effects on pheromone detection vary according to moth species and VPCs. Heptanal, a major component of the floral aroma of linden, activates Phe-ORNs of the noctuid moth *Agrotis ipsilon* (Rouyar, Deisig et al. 2015). Linalool decreases the responses of Phe-ORNs to pheromone compounds in *Spodoptera littoralis* (Party, Hanot et al. 2009). However, linalool and (Z)-3-hexenol synergize the responses of the Phe-ORNs of *Heliothis zea* to its pheromone (Ochieng, Park et al. 2002). A background of VPCs also modulates the firing of MGC neurons masking the response to pheromone (Dupuy, Rouyar et al. 2017). Activity maps obtained by calcium imaging revealed intense activity in the MGC in response to VPCs and various modes of interactions between pheromone and VPCs when they are presented together (Deisig, Kropf et al. 2012). Interestingly, heptanal modified the multiphasic response-pattern of MGC-neurons to pheromone, decreased the response, but improved their capacity to encode pulsed stimuli (Chaffiol, Kropf et al. 2012, Chaffiol, Dupuy et al. 2014). There is now evidence that at least some interactions take place at the OR level. Competitive fluorescence binding assays confirmed that plant odorants compete with the natural pheromone component, Z11 hexadecenal, for binding on HR13, a pheromone receptor of *Heliothis virescens* males (Pregitzer, Schubert et al. 2012). A sex pheromone receptor of the codling moth, *Cydia pomonella*, also binds the plant volatile pear ester (Bengtsson, Gonzalez et al. 2014).

The ecological importance of VPC-pheromone interactions in natural conditions was recently questioned (Badeke, Haverkamp et al. 2016) because, although pheromone attraction of *H. virescens* males was significantly impaired in a concentration-dependent manner after adding single VPCs, their pheromone-guided flight behavior was not influenced by the natural emissions of two host-plants in a wind tunnel. Badeke et coll concluded that the pheromone-VPC interactions only occur at supra-natural concentrations of VPCs. However, insect olfactory communication is now challenged by very fast and intense changes in VPC concentrations caused by human activities. Increasing atmospheric concentrations of O_3_ and other oxidants affect the life-time of odorant molecules in the atmosphere (Atkinson and Arey 2003) with concern for the biotic interactions involving exchange of chemical signals such as pollination (Fuentes, Roulston et al. 2013, Fuentes, Chamecki et al. 2016). Furthermore, plant metabolism is sensitive to global warming and increasing concentrations of CO_2_ and O_3_, which modifies the amounts of VPCs they release in the atmosphere (Lathière, Hewitt et al. 2010, Penuelas and Staudt 2010). Due to these changes in the olfactory landscapes in which insects live it becomes crucial to better understand the impact of olfactory background on insect communication. Pheromone-VPC interactions are a good model to address these questions.

The present research aimed to further investigate the effects of a VPC background on pheromone perception in the moth *A. ipsilon*. We used extracellular electrophysiology and calcium imaging to measure the responses of Phe-ORNs and MGC-neurons to the sex pheromone in simple or composite backgrounds of VPCs. Maps of activities were built using calcium imaging to visualize which areas in moth antennal lobes were affected by VPCs. To stimulate the moth antennae, we used a protocol that mimics the expected natural olfactory landscape in which a pheromone puff must be detected against an odor background (Rouyar, Deisig et al. 2015, Dupuy, Rouyar et al. 2017). Accordingly, short pheromone puffs were delivered over longer-lasting VPC backgrounds. We chose a panel of VPCs with different chemical structures and physicochemical properties representative of the odorant variety encountered by a moth searching for a mate. We first clarified the dose-response effects using single VPCs to create the background. A significant effort was paid to improve the control of stimulus intensity and to establish dose-response relationships. We evaluated the intrinsic activity for each VPC and determined their type of interaction with pheromone. We compared impact of VPC background at antenna and antennal lobe levels. Then, we prepared diverse binary, ternary and quaternary blends of VPCs to investigate interactions between VPCs and to determine whether properties of the blend could be deduced from the properties of single VPCs. Our data confirms that common VPCs interfere with the moth pheromone system in a dose-dependent manner. Activity varies among VPCs. The activity of a blend reproduces that of the most active component with only few interactions between components. Our data will contribute to improve prediction of the impacts of global changes on pheromone communication in insects and exchanges of chemical information within ecosystems.

## Material and methods

### Insects

Four-day-old *A. ipsilon* adult males were obtained from a laboratory population. Larvae were fed on an artificial diet. Males were separated from females at the pupal stage to ensure that the males used for the experiments had never been in contact with the pheromone. Emerging adults were collected every day, kept under a 16h day 8h dark photoperiod and fed with sucrose solution *ad libidum* until the day of the experiment. All experiments were performed during the scotophase.

### Chemicals

The main constituent of the sex pheromone blend of female *A. ipsilon*, (Z)-7-dodecenyl acetate (Z7-12:Ac; CAS 14959-86-5), was purchased from Pherobank (purity > 99%). Based on the literature, we selected eight VPCs with different chemical structures and physicochemical properties to be representative of the variety of odorants that can be encountered by a male moth searching for a mate in an agricultural landscape in mainland France. The monoterpene linalool, the aromatic heterocyclic indole, and the bicyclic sesquiterpene β-caryophyllene have been identified in constitutive and herbivore-induced emissions of *Zea mais*, one of the host crop plants of *A. ipsilon* (Degen, Dillmann et al. 2004). α-pinene is a common monoterpene, emitted by oak and other perennial species that grow on field edges (Staudt and Bertin 1998). The unsaturated hydrocarbon isoprene is one of the most abundant VPC in the atmosphere (Kesselmeier and Staudt 1999); it is released among others by poplars planted to create windbreak hedges or for production wood (Schnitzler, Louis et al. 2010). (Z)-3-hexenyl acetate and (E)-2-hexenal are green leaf volatiles (GLVs) produced by numerous plant species in response to biotic or abiotic stress (Holopainen and Blande 2013). Eucalyptol is a cyclic monoterpene released by flowering weeds, among which *Artemisia annua*, a common weed in maize fields (Jerkovic, Mastelic et al. 2003). VPCs were diluted in light mineral oil (CAS 8042-47-5). VPC synthetic standards (Table S1) and mineral oil were obtained from Sigma Aldrich.

### Odor stimulus delivery

The stimulus delivering device consisted of an ensemble of microvalves enabling to deliver VPCs from separate sources in the main air stream of a glass tube (length 200 mm, inner diameter 9 mm) whose distal end was positioned 20 mm from the insect antenna, while the pheromone stimuli were delivered through a lateral input (Figure S1). Air was charcoal-filtered and humidified.

Pheromone stimuli were delivered as air puff (167 mL/min) through a Pasteur pipette containing a piece of filter paper loaded with 10 ng (unless mentioned) of Z7-12:Ac diluted in 1 μL of hexane. Hexane was left to evaporate for 30 s before inserting the pheromone-loaded paper into the pipette. Air passage through the pipette was commanded by an electrovalve (LHDA1233215H, The Lee Company, Westbrook, USA). The pipette tip was inserted into a hole on the side of the glass tube, 150 mm upstream of its distal end.

VPC sources consisted of 4 mL glass vials containing 1 mL of a single VPC diluted in mineral oil, or mineral oil only (control). The air stream was divided into 8 parallel flows (200 mL/min each) with an airflow divider (LFMX0510528B, The Lee Company), each of which directed towards a 3-way electrovalve (LHDA1223111H, The Lee Company). Normally opened (NO, non-odorized) and normally closed (NC, odorized) exits of the eight valves were connected either to empty vials or to VPC source vials, respectively. All outlets of odorized and non-odorized vials were connected to the proximal end of the glass tube. Thus, valve opening did not modify the total airflow received by the antenna (1.6 L/min). All tubing downstream from the valves was made of Teflon (internal diameter 1.32 mm). Vials were connected to the tubing with stainless steel hypodermic needles inserted through a Teflon septum. For delivering single VPCs, the Teflon tubes at vial outlet were directly connected to the main glass tube (Figure S1A). VPC mixing was achieved by opening several valves simultaneously and mixing odorized airflows in a low dead-volume manifold (MPP-8, Warner Instruments, Figure S1B). For each of the 8 valves, the NO and the NC exits were connected together before entering one of the 8 manifold inlets.

### Electrophysiology

For single sensillum recordings, male moths were briefly anesthetized with CO_2_ and restrained in a Styrofoam holder. One antenna was immobilized with adhesive tape. A tungsten electrode was inserted into the antenna to serve as a reference. We targeted the Phe-ORNs tuned to Z7-12:Ac. The recording electrode was therefore inserted at the base of a long trichoid sensillum located along antennal branches.

For extracellular recordings from MGC-neurons, male moths were restrained in a cut pipette tip, leaving the head exposed, and immobilized with dental wax. The head capsule was opened, and the brain exposed by removing all muscles and mouthparts above it. The neurolemma was carefully removed from the antennal lobe in order to allow electrode penetration. The recording electrode was made from a glass micropipette whose tip was manually broken to a diameter of 2 μm and filled with (in mM): NaCl 150, KCl 4, CaCl_2_ 6, MgCl_2_ 2, Hepes 10, Glucose 5 (pH 7.2, osmotic pressure 360 mOsm/L adjusted with mannitol). The preparation was constantly perfused with the same solution once the brain capsule was opened. The reference electrode was a silver wire inserted at the rear of the head capsule in contact with brain tissues. The recording electrode was slowly inserted inside the MGC until the appearance of a clear single-unit firing activity. Extracellular recordings from *A*. *ipsilon* AL sample only neurons with a large neurite (Rospars, Gremiaux et al. 2014) so we expected to record mainly projection neurons (PNs) rather that local interneurons (LNs). Recordings were done using a CyberAmp 320 controlled by pCLAMP10 (Molecular Devices, San Jose, USA). The biological signal was amplified (×2000), band-pass filtered (1-3000 Hz) and sampled at 10 kHz with a Digidata 1440A acquisition board (Molecular Devices). Spikes were sorted using Spike 2 software (CED, Oxford, Great Britain).

### Calcium imaging in antennal lobes

Male moths were restrained in a Plexiglas chamber and the head was fixed. The head was opened and muscles and mouthparts removed to gain access to the brain. Then, 20 μL of a dye solution (50 μg Calcium Green 2-AM dissolved with 50 mL Pluronic F-127, 20% in dimethylsulfoxide, Molecular Probes, Eugene, OR, USA) was bath-applied for at least 1 hour, before being washed with Ringer. For recordings a T.I.L.L. Photonics imaging system (Martinsried, Germany) was coupled to an epifluorescent microscope (BX-51WI, Olympus, Hamburg, Germany) equipped with a 10× (NA 0.3) water immersion objective. Signals were recorded using a 640 × 480 pixel 12-bit monochrome CCD camera (T.I.L.L. Imago, cooled to −12°C). The acquisition rate was set at 5 frames/s with an acquisition time of 15 ms. Identification of individual glomeruli was done by superposing activity maps using Adobe Photoshop (Version CS2). The order in which the background VPCs were tested was kept constant in the case of calcium imaging and stimuli with VPCs as a background to the pheromone were alternated with stimuli with VPCs alone.

Raw data analysis was done using custom–made software written in IDL (Research Systems Inc., Colorado, USA) and Visual Basic (Microsoft Excel). After noise filtering and bleaching correction, relative fluorescence changes (δF/F) were calculated as (F-F_0_)/F_0_ (where F_0_= reference background). For each glomerulus, the time course of δF/F was calculated by averaging 25 pixels (5 × 5) at the center of each glomerulus.

### Stimulation sequences

First, we evaluated the effects of single VPCs on the activity of MGC-neurons and Phe-ORNs and determined their active doses. Stimulation sequences consisted of a short pheromone puff delivered in the middle of a 5 s VPC background presentation. Linalool, whose ability to activate the MGC-neurons of *A. ipsilon* had been already demonstrated (Dupuy, Rouyar et al. 2017), was used to establish the dose used for all the VPCs of the panel. We first measured the activity of different dilutions of linalool on MGC-neurons, showing that a dilution of 1% triggered a clear firing activity. We adjusted dilutions of the other VPCs to deliver similar concentrations in the stimulus flow in spite of the differences of volatility according to procedures proposed by Munch et al (Munch, Schmeichel et al. 2013) based on data from (Cometto-Muniz, Cain et al. 2003) (Table S1). For electrophysiological recordings, the pheromone puff lasted 200 ms and started 2.8 s after background onset (Figure S2). For calcium imaging, the slow response dynamics of the fluorescence signal required to adapt stimulus duration. The pheromone puff lasted 1 s and started 2 s after background onset. Successive stimuli on the same preparation were separated by 30 s (antennal lobe recordings) or 60 s (single sensillum recordings and calcium imaging). Valve opening/closing sequences were computer-controlled with millisecond accuracy. The VPC were presented in random order, except for calcium imaging. In imaging, stimuli with VPCs as a background to the pheromone were alternated with stimuli with VPCs alone.

To measure the effects of binary blends of VPCs, binary mixtures and their constituents alone were tested at 4 concentrations each, while the pheromone dose was kept constant. In order to control precisely the ratios in the background, we compensated differences in volatility among the VPCs to be blended. To this end, we measured their air-mineral oil partition coefficients (Khl, Table S1) and used them to calculate the mineral oil concentration (C_l_) necessary to obtain the desired concentration in the headspace (C_h_) of a closed, equilibrated source:

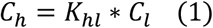

Partition coefficients K_hl_ were measured by injecting the headspace of closed equilibrated sources containing known concentrations of VOCs into a calibrated GC-FID (Conchou et al, in preparation). When an airflow is run through such a source, the concentration in the headspace drops to a fraction of the initial concentration, then remains stable until the source starts to exhaust. For the VPCs included in the blend experiments, we found that this fraction is independent of odorant identity (Conchou et al., in preparation). Therefore, odorant concentration delivered from a source is related to the initial concentration inside the closed equilibrated source by a dilution factor that does not depend on the odorant. Assuming that this proportionality remains true for the concentrations delivered on the antenna (i.e. no bias induced by differential adsorption on tubing walls), we expressed all delivered VPC concentrations relative to an arbitrary unit (AU) where one AU was defined as the molar concentration delivered on the antenna from a source containing 1% linalool in mineral oil.

Binary blends were obtained by opening simultaneously the valves controlling the airflow through the sources containing the single VPCs to be blended while closing a compensation flow. Therefore, if the concentrations of blend components, when tested separately, are noted A and B, the blend was tested at the concentration A + B. The order of presentation of individual VPCs and of their combination was randomized. For each background type, 4 successive stimuli were presented at increasing concentration. A pheromone puff with no background was presented as a control at the beginning of the recording and after the highest concentration of each background type. Stimulus intensity was checked repeatedly with a photoionization detector (PID, from Aurora Scientific Inc, Aurora, Canada). Experiments began by ejecting a VPC stimulus from all sources away from the recorded antenna because PID measurements showed that the first stimulus was more intense than the following ones, while subsequent stimuli from the same VPC source had the same intensity if 5 to 10 minutes apart. Then, the glass tube was focalized on the antenna and intervals between successive openings of the same VPC valve were always 5 or 10 minutes.

Finally, we tested blends of 3 or 4 VPCs as backgrounds to the pheromone stimulus. The background and pheromone concentrations were kept constant. We alternated presentation of these more complex backgrounds with backgrounds consisting in a single VPC or a binary blend of its components. Each of these 2 composite backgrounds and the corresponding simple backgrounds were presented twice to each neuron in random order. Control stimuli (pheromone without background) were presented at the beginning of each recording and after each pair of composite and simple background stimuli.

### Electrophysiology data processing and statistical analysis

Analysis of firing activity was performed using custom programs developed under R (R Core Team 2013). An instantaneous firing rate metric (Blejec 2005) was used to draw peri-stimulus firing curves. For each individual recording, a firing rate was calculated for every spike using the two preceding and two following spikes. Then, we calculated the average firing rates over successive 75 ms long time bin in each recording. Individual neuron firing rates/bin were finally averaged over all sampled neurons to draw peri-stimulus curves.

We calculated the maximum firing frequency within four time-windows (TWs) covering the successive phases of the two stimuli. The limits of each TW are the valve opening times shifted to consider the travel time of the odorized airflows from the valves to the antenna (Figure S2). TW1 (from 0 to 5.2 s) covered the period before background application and was used to measure the spontaneous activity. The phase corresponding to VPC background onset until pheromone puff was split into two TWs. TW2 (5.2-7.0 s) corresponds to the fast rise in the neuron firing that followed the background onset; TW3 (7-8 s) covered the period during which the firing decreased notably compared to the peak, but stayed significantly above the spontaneous activity for the most active PVCs; TW4 (8-8.5) covered the response to pheromone. To correct for differences in spontaneous activity between neurons, we calculated the **response to background** by subtracting the mean firing frequencies in TW1 (mean_TW1_) from the maximum firing frequency reached in TW2. Similarly, the **pheromone response** was calculated by subtracting mean_TW1_ from the maximum firing frequency reached within TW4 (max_TW4_). We estimated the capacity of neurons to extract the pheromone signal from the background, the **pheromone salience,** by subtracting pre-pheromone activity level in TW3 from max_TW4_.

For experiments evaluating the effects of single VPCs we used pairwise paired t tests on all possible pairs of background types to compare values of response to background, response to pheromone and pheromone salience in the different backgrounds. False discovery rate was controlled using the Benjamin-Hochsberg’s procedure (FDR < 0.05).

For experiments comparing blends to their constituents we used a permutational MANOVA (PERMANOVA, function adonis() under R package vegan (Anderson 2001, Jari Oksanen, Blanchet et al. 2019) to evaluate how background composition and dose affected neuronal responses, the three measured variables taken together. The PERMANOVA used a Euclidian distance matrix calculated from response to background, response to pheromone, and pheromone salience. For significance testing, permutations were restricted within individual recordings (parameter “strata”) to account for the non-independence of observations made on the same neuron. Whenever significant differences were found, we further tested differences between the blend and each constituent using pairwise PERMANOVAs between all three pairs of background types. False discovery rate was controlled using the Benjamin-Hochsberg’s procedure (FDR < 0.05). Note that under this analysis, dose was considered a categorial variable, which may not be sufficient to accurately describe the way neurons respond to a blend of two agonistic components. Therefore, for the blend (Z)-3-hexenyl acetate / linalool we further refined the analysis by taking the actual concentrations into account through modeling the dose-response curves. Dose-responses to odorants are usually modelled using Hill’s equation (Rospars, Lansky et al. 2008): the response is described as a function of ligand concentration (C), and depends on a maximal response intensity (R_max_), a concentration at half maximum (EC_50_) and Hill’s coefficient (n):

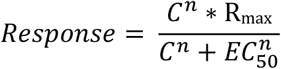

Our response to background data were not appropriate for a fit of Hill’s equation because the saturation was never reached making it impossible to estimate R_max_. However, in the case of pheromone response and of pheromone salience, the aim was to model the decrease in response intensity observed in the presence of an agonist background. For that purpose, response intensity or salience in the absence of background could be considered as R_max_. Since individual neurons differed in their responsiveness to the pheromone, we normalized all observed pheromone response and salience values to the considered neuron’s corresponding R_max_:

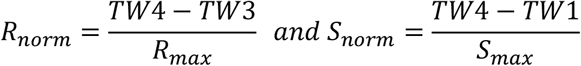

This also allowed to simplify Hill’s equation by setting R_max_ = 1. Subtracting this simplified Hill’s equation from 1 produces a curve that decreases with increasing C, which is empirically appropriate for fitting on the observed decreasing response to pheromone or pheromone salience as function of the background dose:

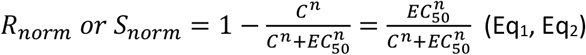

We fitted equations Eq1 and Eq2 using the non-linear regression function nls2() (R package nls2, G. Grothendieck, 2013, CRAN deposit) to estimate EC_50_ and n for the Phe-ORN recordings pooled together (n = 16), and for each background type separately, using total background concentration in arbitrary units as C. We compared the dose-responses observed under blended background and under single constituent’s background by checking whether the confidence intervals for the fitted parameters did or did not overlap.

## Results

### Single VPCs elicit calcium responses in ordinary glomeruli and in the MGC

All tested VPCs induced a calcium response (Ca-response) in some parts of the antennal lobes. Eight areas showing increase in fluorescence in response to the presentation of at least one VPC were putatively identified as distinct ordinary glomeruli (OG). Activity patterns were very different among VPCs (Figure 1), with some VPCs activating repeatedly more than half of the region of interest identified as putative OGs ((E)-2-hexenal, (Z)-3-hexenyl acetate, linalool), while others (β-caryophyllene, eucalyptol and indole) showed more limited activity maps. For α-pinene, an increase in fluorescence in one OG (OG5, Figure 1) was observed in only two preparations. We chose to not include isoprene in calcium imaging experiments due to the limited number of channels available in the odor stimulation device. The pheromone compound, Z7-12:Ac, did not activate any of these OGs.

**Figure 1:**
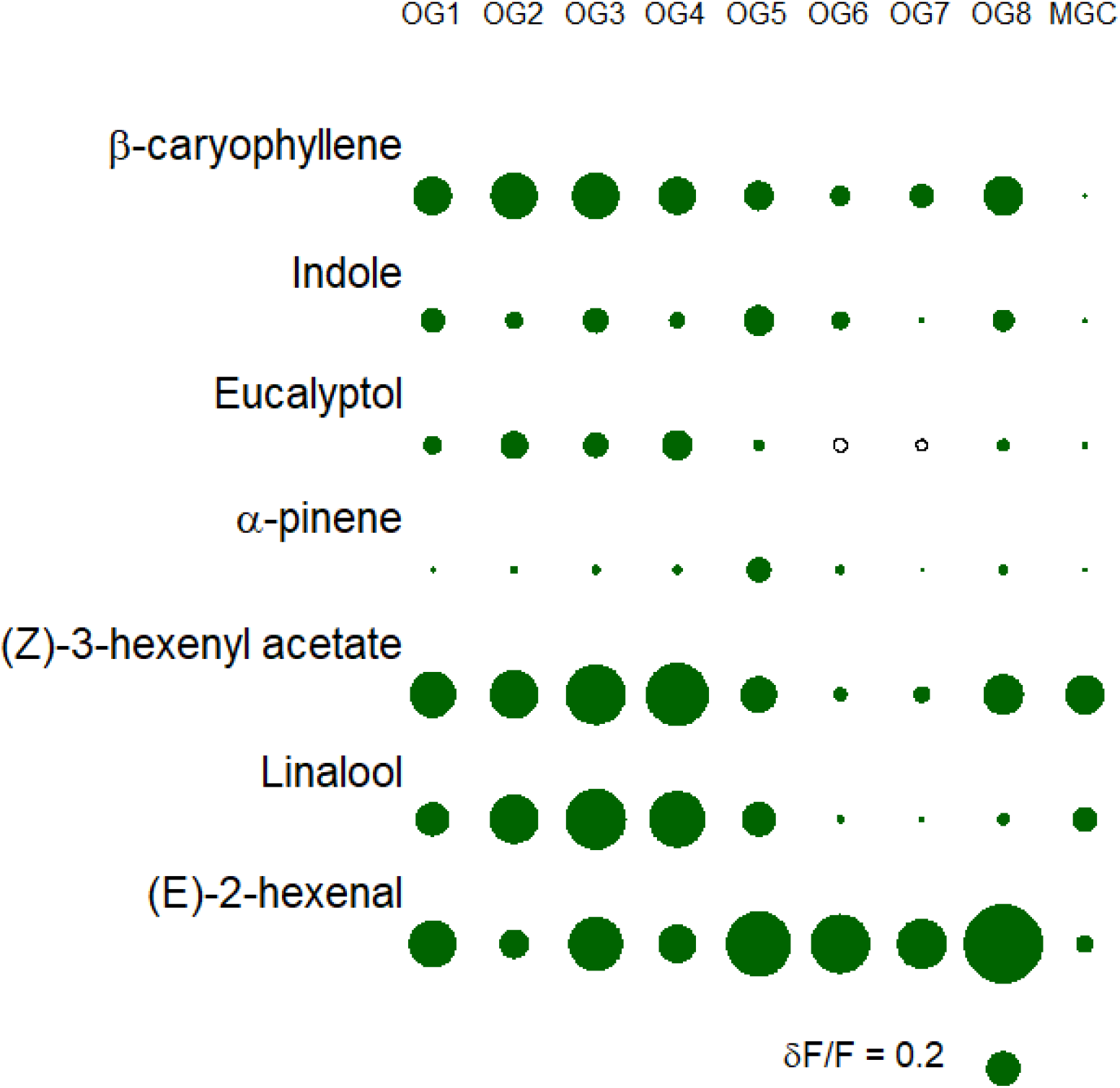
Map of the calcium activity in *A. ipsilon* antennal lobe in response to seven VPCs in different putative ordinary glomeruli (OG1 to OG8) and in the MGC. The diameter of the circles is proportional to the relative fluorescence change (δF/F) calculated by averaging a 25-pixel square at the center of the ROI attributed to each glomerulus (N = 15).

Z7-12:Ac activated a large area in the AL that we identified as the MGC based on previous work in the same insect species (Barrozo, Jarriault et al. 2011, Deisig, Kropf et al. 2012). (Z)-3-hexenyl acetate, linalool and (E)-2-hexenal also triggered a strong Ca-response within the MGC (Figure 1). The Ca-responses to pheromone and VPCs were tonic, lasting the time of the odorant presentation (Figure 2). Eucalyptol, indole, caryophyllene and α-pinene did not evoke significant Ca-response in the MGC.

**Figure 2:**
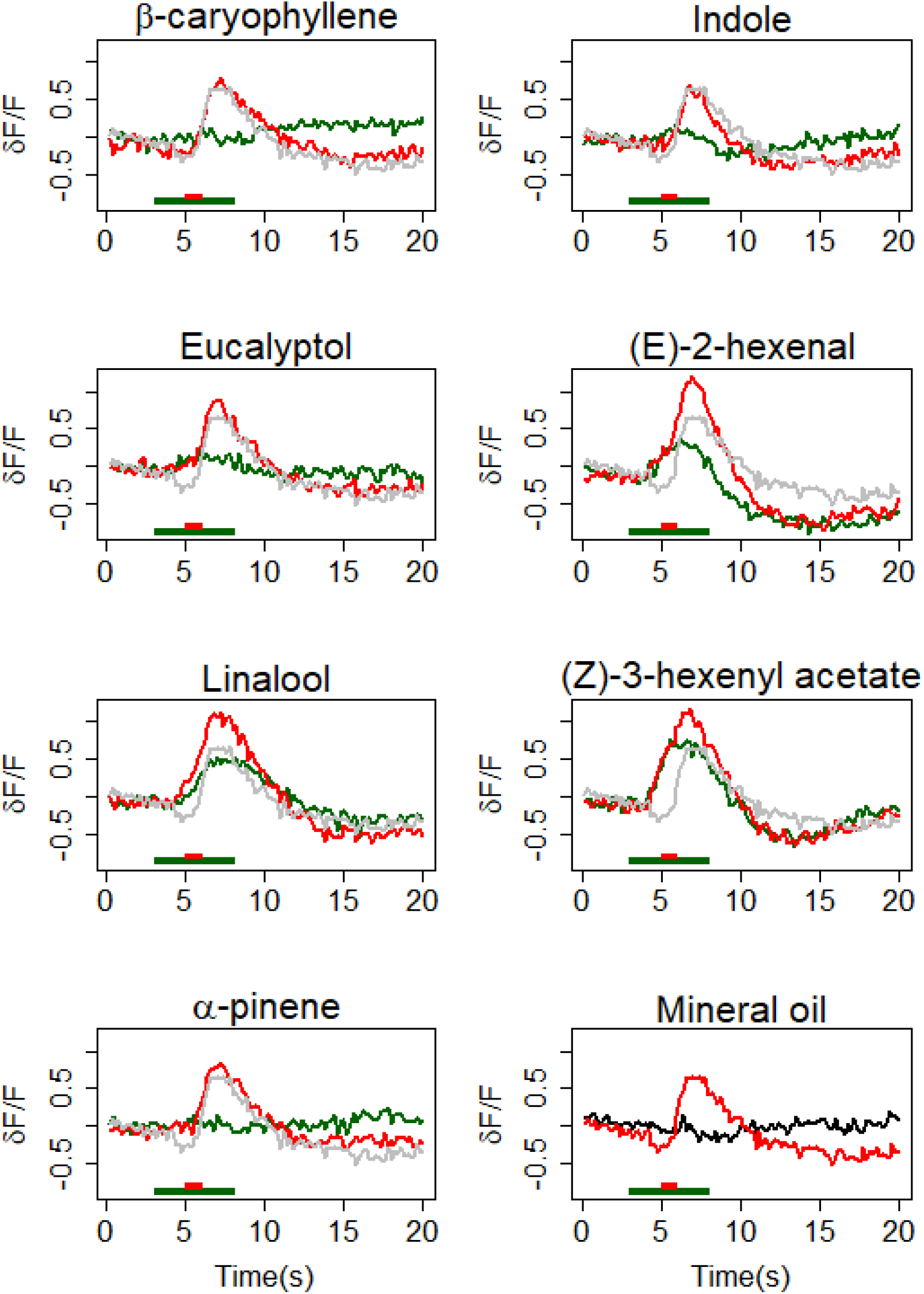
Effects of a VPC background on the calcium-response to pheromone in the MGC area of the antennal lobe. Average time course (N = 15) of δF/F in response to VPCs (green) and to the pheromone in a background of the same VPCs (red) and to pheromone in control background (grey). The last panel presents the responses to the pheromone in the control background (mineral oil only, red) and to the control background (black). Green and red bars at the bottom of each graph mark the VPC and pheromone stimuli, respectively.

When the pheromone was delivered during the VPC-background, Ca-responses to VPC and to pheromone merged to form a single fluorescence-peak (Figure 2). Thus, it was not possible to quantify separately the contribution of the VPC and that of pheromone to the Ca-response. However, the Ca-response to VPC plus pheromone was always significantly larger than that of VPC alone, whatever the VPC (Table 1, test VPC vs VPC + Phe). It was significantly larger compared to pheromone in mineral oil for 2-hexenal, linalool and Z3-hexenyl acetate, the three VPCs which triggered a Ca-response within the MGC (Table 1). For the other four VPCs of the panel, the Ca-response to pheromone in a VPC background was not different from that to pheromone in the control background. The response to pheromone alone was always above that to the VPC alone, even if for some VPCs the difference was not significant (Table 1).

**Table 1:**
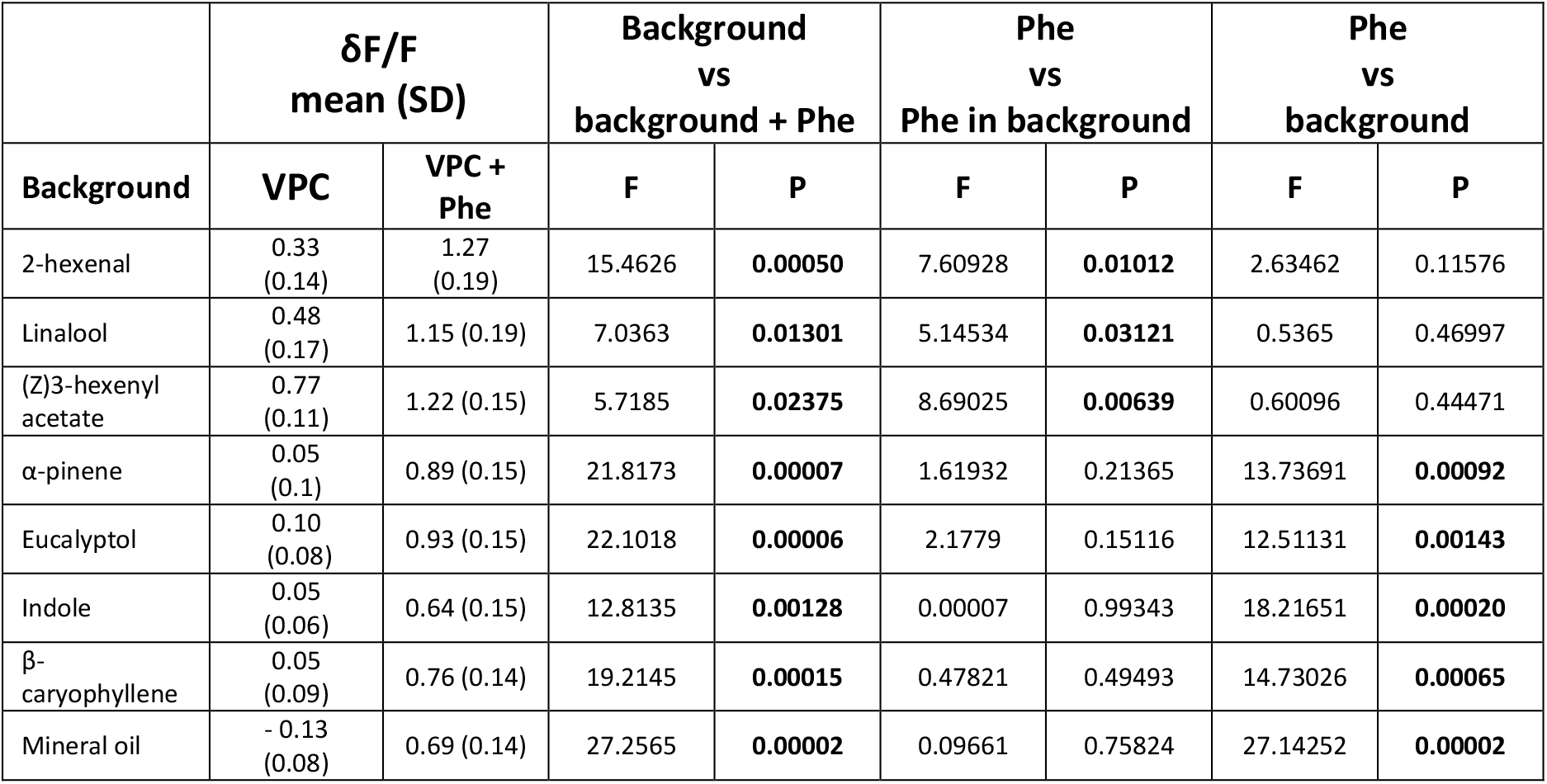
Intensities of calcium responses in the MGC following a 5 s presentation of a background of single VPC or of VPC-background plus 1 s puff of the pheromone component Z7-12:Ac. Means (N = 15) and standard deviations (in brackets) of the maximum δF/F. Statistical tests: One-way ANOVA, 1 degree of freedom.

### Some single VPCs activate MGC-neuron firing and affect their responses to the pheromone

We first tested the impact of 7 doses of linalool (from 0.00001 to 10% in mineral oil) on the firing activity and responses to pheromone in 5 MGC-neurons. Linalool activated the firing of the MGC-neurons in a dose-dependent manner (ANOVA, P < 0.0001) with a threshold at 1% (posthoc paired t-tests, compared to control: response to background P= 0.0631; response to pheromone, P = 0.0019; pheromone salience, P < 0.0001). Recordings showed a fast rise of the firing at the background onset, followed by a plateau lasting the time of the linalool presentation. Linalool 1% was chosen as a reference stimulus for further experiments.

Then, we stimulated the moth antennae with the other VPCs of the panel, adjusting their concentrations in the source vial according to their volatilities to deliver the same VPC quantities as with 1% linalool (see Material and Methods, Table S1). Response varied according to the VPC (ANOVA, P > 0.00001). (Z)-3-hexenyl acetate strongly stimulated the MGC-neurons (Figure 3A; posthoc t-test, response to background: P < 0.0001). Compared to linalool, responses to (Z)-3-hexenyl acetate were more dynamic, showing a fast initial peak with short latency (Figure 3B), followed by a decrease in firing and a sustained plateau until background offset. Increases in firing were also observed in some recordings in response to 0.1% (E)-2-hexenal (Figure 3A-C), but the difference with the control background is not significant (response to background: P = 0.0579) due to the variability of responsiveness of MGC-neurons to this VPC (Figure 3C). Several MGC-neurons showed a decrease in their firing activity at 1% eucalyptol presentation (Figure 3C) suggesting an inhibitory activity but the difference with the control was not found globally significant (response to background: P = 0.0928).

**Figure 3:**
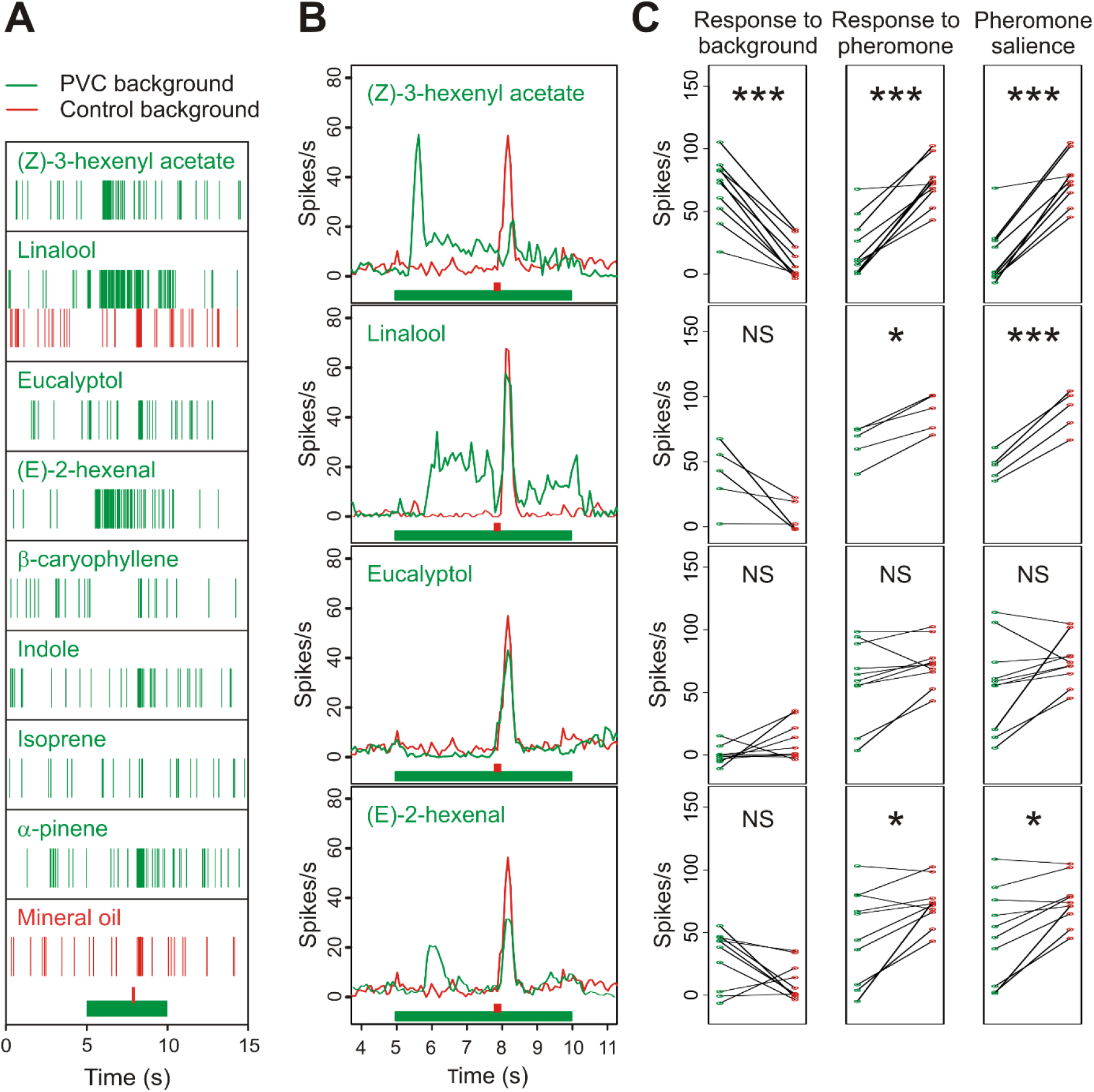
Effects of a VPC background on the firing activity of MGC-neurons and their responses to the pheromone. (A): raster plots of typical individual extracellular recordings. (B): Mean frequency plots showing the fast firing peak at background onset, followed by a sustained firing activity, and their effects on the amplitude of the firing peak at pheromone presentation. In A and B, green and red bars indicate background and pheromone stimuli, respectively. (C): Strip charts comparing individual firing activities in each VPC background (green dots) with the control background (red dots). Firing frequency was measured on appropriate time windows to evaluate: response to background (left column), response to pheromone (middle column), and pheromone salience (right column). * and *** indicate p-values of the paired t test below FDR threshold; NS = p-value above FDR threshold. N = 5 for linalool and 10 for other compounds.

Contrary to the Ca-imaging experiments, it was possible to isolate the increase in firing activity in response to the pheromone puff and to measure the pheromone salience. Response to pheromone was dramatically reduced in a (Z)-3-hexenyl acetate background (P < 0.0001), indicating the occurrence of mixture suppression and resulting in an almost complete masking of the pheromone (Figure 3C). Reduction of the response to pheromone was also observed in presence of the (E)-2-hexenal background (P = 0.0169). The combination of response to background and reduction in response to pheromone resulted in a significant reduction of pheromone salience in the presence of (E)-2-hexenal (P= 0.0061), linalool (P = 0.0003) or (Z)-3-hexenyl acetate (P < 0.0001) backgrounds.

Although the firing activity seemed lower in the presence of eucalyptol in some MGC-neurons (Figure 3C), this decrease was not significant (P = 0.0579) and response to pheromone and pheromone salience were not statically different between eucalyptol and control background groups. β-caryophyllene, indole, isoprene and α-pinene neither elicited significant responses to background nor significantly altered the response to pheromone (Figure S3).

### Single VPCs modulate the Phe-ORN spontaneous firing and affect their responses to pheromone

We then recorded the responses of Phe-ORNs to pheromone in the presence of the same VPCs at the same concentrations as for MGC-neurons and compared the effects of VPC backgrounds on Phe-ORNs and MGC-neurons to clarify whether interactions took place at peripheral or AL levels.

The impact of the (Z)-3-hexenyl acetate background on ORNs was similar to its effect on MGC-neurons (Figure 4). Phe-ORNs responded to the background with a significant increase in firing (P < 0.0001). However, compared to MGC-neurons, their firing activity decreased more slowly with a longer peak tail (Figure 4B). The response to the pheromone was significantly reduced (P = 0.0004). The firing peak in response to the pheromone was hardly visible (Figure 4B) and accordingly pheromone salience was strongly decreased (P < 0.0001) (Figure 4C).

**Figure 4:**
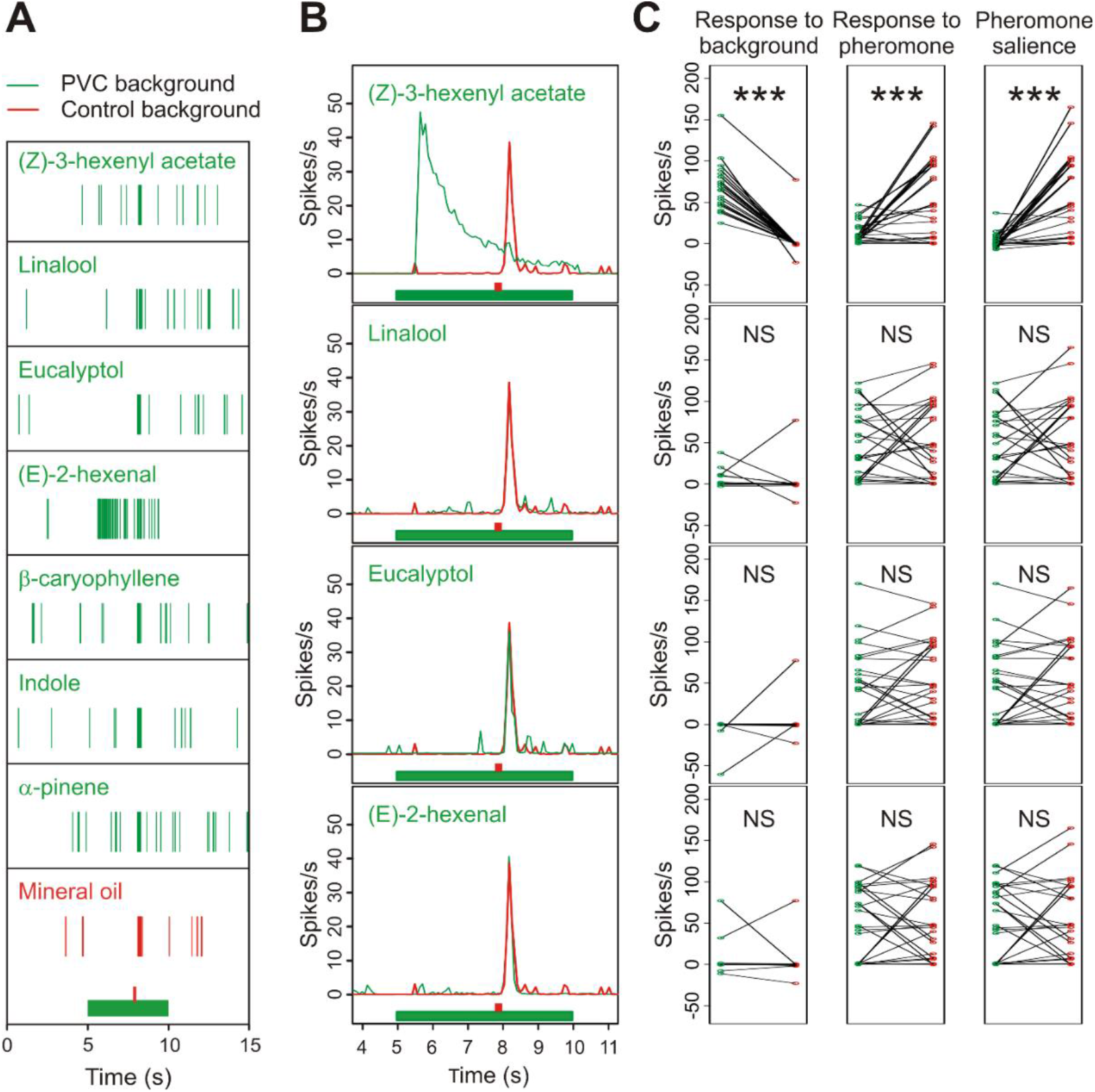
A background of (Z)-3-hexenyl acetate, linalool, eucalyptol or (E)-2-hexenal modifies the firing activity of Phe-ORNs and their responses to pheromone. (A): raster plots of samples of single sensillum recordings. (B): Mean frequency plots showing the time-course of the firing in response to background presentation and pheromone pulse. In A and B, green and red bars indicate background and pheromone stimuli, respectively. (C): Strip charts comparing individual neuron firing activities in each VPC background (green dots) with the control background (red dots). Firing frequency was measured on appropriate time-windows to evaluate the response to background (left column), response to pheromone (middle column), and pheromone salience (right column). N = 26. Stars indicate p-values of the paired t test below FDR threshold; NS = p-value above FDR threshold. N = 5 for linalool and 10 for other compounds.

Surprisingly considering their activity on MGC neurons, Phe-ORNs responded neither to a 1% linalool background (Figure 4B; P = 0.6724) nor to a 0.1% (E)-2-hexenal background (P= 0.6709). Response to pheromone and pheromone salience were not altered by linalool (P = 0.5710 and 0.4598, respectively) or (E)-2-hexenal (P = 0.9933 and 0.9883, respectively).

In agreement with its lack of activity in MGC-neurons, no significant effect of α-pinene was observed in Phe-ORNs, either for response to background (Figure S4; P = 0.5447), response to pheromone (P = 0.6244), or pheromone salience (P = 0.8908). The indole background did not activate Phe-ORNs (Figures 4 and S4; response to background P = 0.73542), and did not modify response to pheromone (P = 0.8275) and pheromone salience (P = 0.1126).

To highlight putative inhibitions, we applied backgrounds as short pulses over a sustained pheromone stimulation (Figure S5). Compared to control, Phe-ORNs stimulated by the pheromone had a lower activity during presentation of the eucalyptol background than before (Figure SI-5; P = 0.023 for the comparison of the difference in number of spikes in TW4 and TW3). Such inhibitory responses were not observed for (E)-2-hexenal (P = 0.0621), α-pinene (P = 0.254), indole (P = 0.9611), β-caryophyllene (P = 0.1920) and linalool (P = 0.8499). We also noted a significant increase for (Z)-3-hexenyl acetate (P = 0.0068), due to the response to the background.

### Binary blends produce the effects of their most active component

We prepared binary blends by combining VPCs with contrasted impacts on the pheromone detection and compared their effects at different doses to that of their individual components. To facilitate comparisons between single components and their blends the concentrations delivered to the antenna were expressed in arbitrary units (AU) as defined in the material and methods.

We first mixed the agonist (Z)-3-hexenyl acetate with α-pinene, which neither stimulated Phe-ORNs nor modified their response to pheromone. To increase the probability of evidencing an interaction between the two PVCs when blended we doubled the proportion of α-pinene relatively to that of (Z)-3-hexenyl acetate (ratio 2:1). The effects of (Z)-3-hexenyl acetate were clearly dose-dependent (Figure 5; global PERMANOVA, dose effect P = 0.001) and responses to the background increased with the concentration of (Z)-3-hexenyl acetate (Figure 5 A and B). The responses to the pheromone were more strongly attenuated at higher concentrations (Figure 5C). Consequently, the pheromone salience decreased when increasing the background concentration (Figure 5D). The firing of Phe-ORNs was the same in a background of α-pinene as in the control background, whatever the α-pinene concentration (Figure 5A and B; pairwise PERMANOVA, α-pinene versus control backgrounds, P = 0.939). When the binary blend was presented as background, neuron activities were not different from those observed under a background of (Z)-3-hexenyl acetate alone (Figure 5; pairwise PERMANOVA on (Z)-3-hexenyl acetate versus blend backgrounds, background type effect P = 0.262).

**Figure 5:**
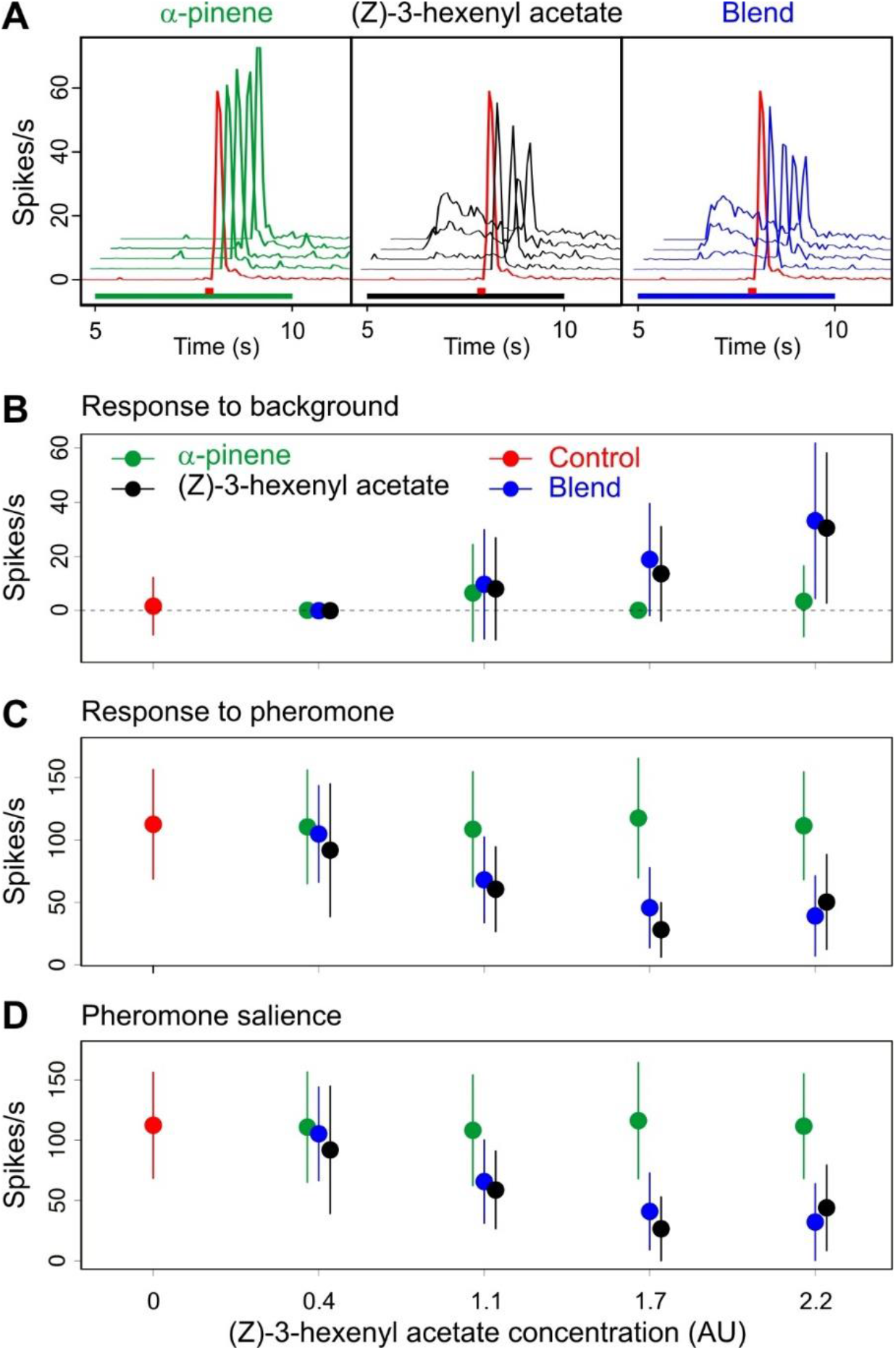
Effects on Phe-ORNs of a blend of α-pinene and (Z)-3-hexenyl acetate at 2:1 ratio as background to pheromone. (A): Mean frequency plots (N = 15) showing the time-course of the firing during background presentation and after pheromone pulse at increasing concentrations of background; below the plots green (α-pinene), black ((Z)-3-hexenyl acetate) or blue (blend) bars indicate background stimulus and red bars indicate pheromone stimulus. (B to D): Effects of background dose and composition on response to background (B), response to pheromone (C) and pheromone salience (D). Means and standard deviations, N = 15.

Next, we tested a blend of linalool, which showed agonist activity, with eucalyptol, a VPC that inhibited some MGC neurons and Phe-ORNs when presented over a sustained pheromone stimulation, at a ratio of 1:2 (Figure 6). At the tested concentrations, eucalyptol alone did not have any noticeable impact on the neuronal activity (Figure 6B; pairwise PERMANOVA eucalyptol vs control backgrounds, background type effect P = 0.212). Linalool showed a dose-dependent agonist activity (global PERMANOVA, background type effect P = 0.001, dose effect P = 0.01): response to the linalool background was apparent only at 4AU (Figure 6B) while the reduction in pheromone response and salience appeared at 2AU (Figure 6C and D). The blended background did not modify neuronal activity any differently from linalool alone (pairwise PERMANOVA linalool versus blended background, background type effect P = 0.570, Figure 6B-D).

**Figure 6:**
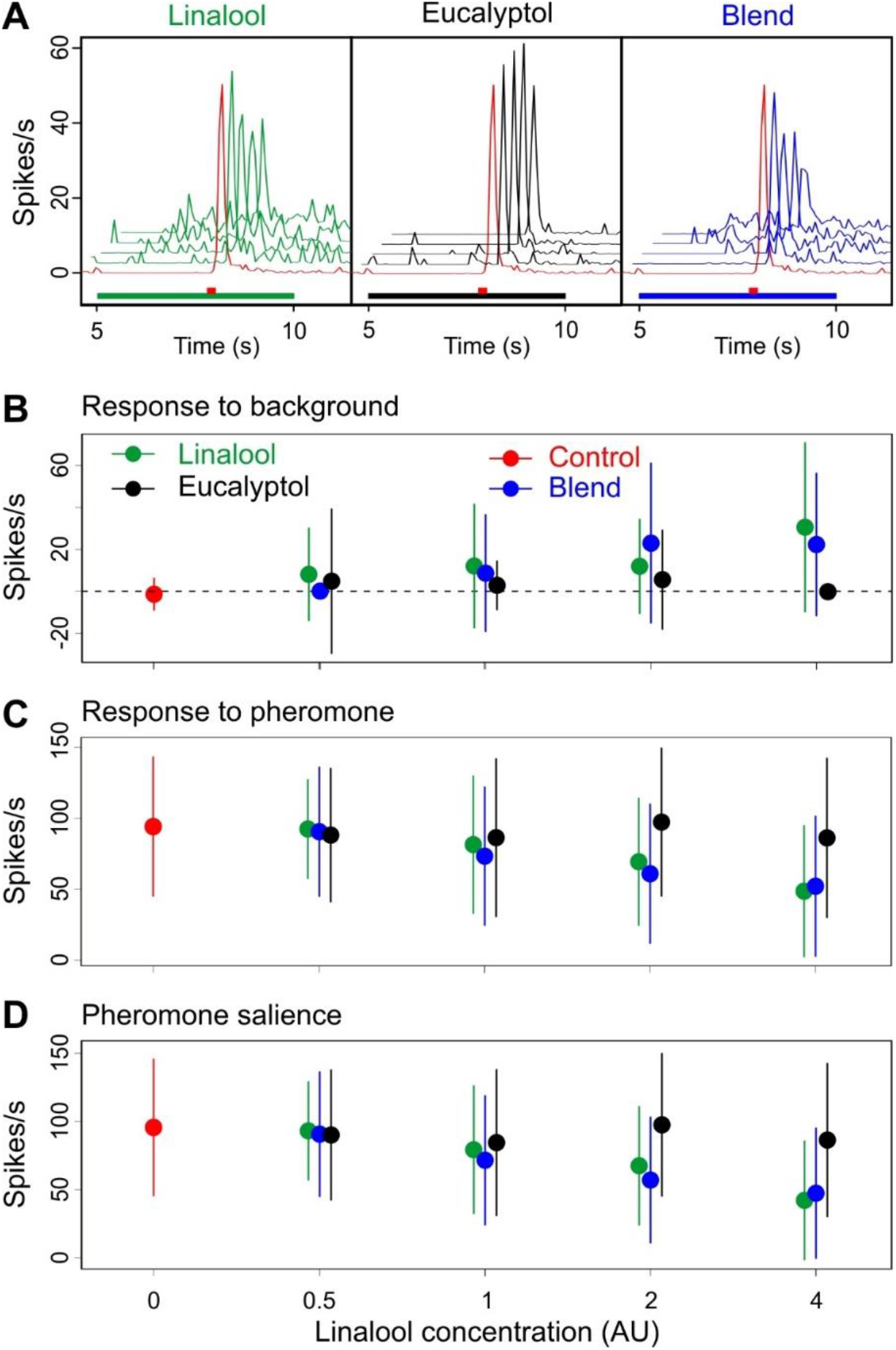
Effects on Phe-ORNs of blending linalool with eucalyptol at a 1:2 ratio as background to a pheromone stimulus. (A): Mean frequency plots (N = 17) showing the time-course of the firing during background presentation and after pheromone pulse; below each plot a green (linalool), black (eucalyptol) or blue (blend) rectangle indicates background presentation and a red rectangle the pheromone stimulus. (B to D): Effects of background dose and composition on response to background (B), response to pheromone (C) and pheromone salience (D). Means and standard deviations, N = 17.

We then mixed the two agonists (Z)-3-hexenyl acetate and linalool at a 1:1 ratio and compared the blend effects to that of its single constituents. Comparison of the activity of Phe-ORNs in response to increasing doses confirmed that linalool was a weaker agonist than (Z)-3-hexenyl acetate (pairwise PERMANOVA (Z)-3-hexenyl acetate versus linalool, background type effect, P = 0.0010; Figure 7A and B). Adding the two VPCs to each other resulted in a significantly stronger activity of the blend compared to linalool alone (pairwise PERMANOVA linalool versus blend, background type effect, P = 0.001). The blend was also more active than (Z)-3-hexenyl acetate alone at the two highest concentrations (pairwise PERMANOVA, (Z)-3-hexenyl acetate versus blend, background type effect P = 0.001). However, since the global blend concentration was the sum of that of its constituents we could not determine which type of blend interaction occurred between (Z)-3-hexenyl acetate and linalool. To this end, we modeled the background doses - response to pheromone and pheromone salience curves for each of the three background types. Our modeling approach revealed that the parameters of the model curves for the blend were always closer to the estimation for (Z)-3-hexenyl acetate than to linalool with stronger estimated EC_50_ value and a lower coefficient (Figure 8 and Table S2) confirming (Z)-3-hexenyl acetate salience in the background effects.

**Figure 7:**
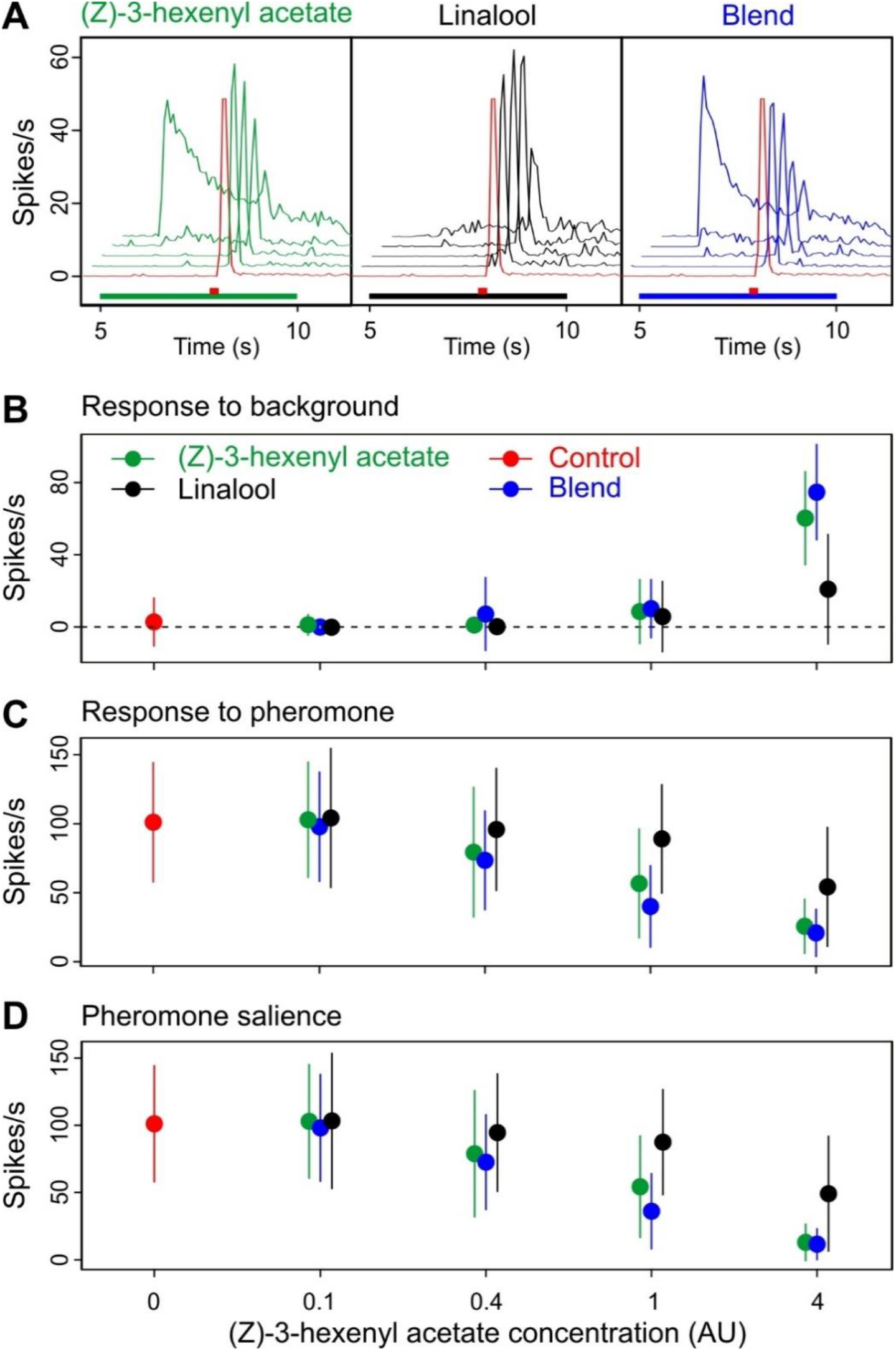
Effects on Phe-ORNs of blending two agonists VPCs, (Z)-3-hexenyl acetate and linalool, at a 1:1 ratio as background. (A): Mean frequency plots showing the time-course of firing during background presentation and after pheromone pulse. Below the plots, green ((Z)-3-hexenyl acetate), black (linalool) or blue (blend) rectangles indicate background presentation; a red rectangle indicates the pheromone stimulus. (B to D): Effects of background dose and composition on response to background (B), response to pheromone (C) and pheromone salience (D). Means and standard deviations, n= 16.

**Figure 8:**
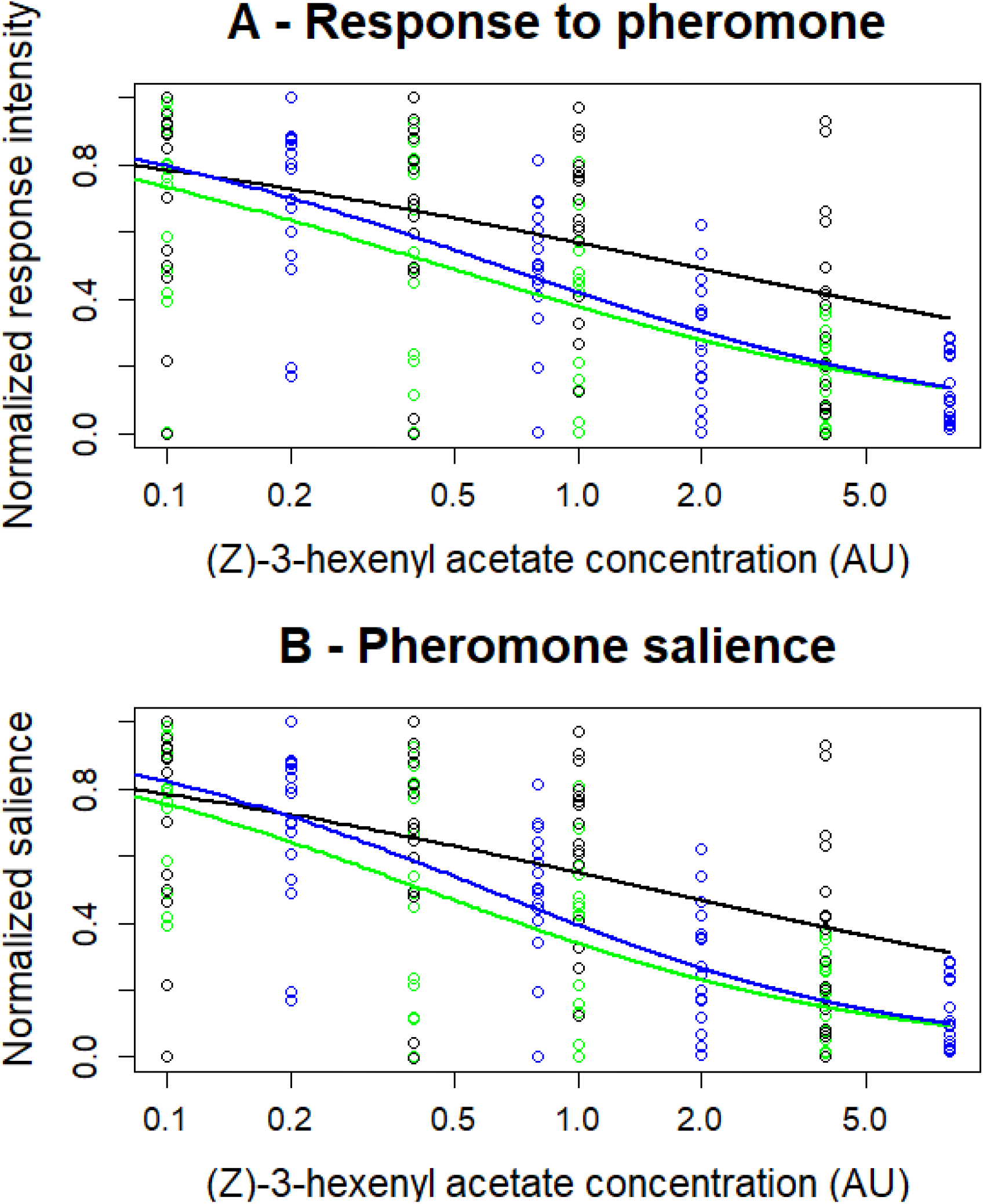
Modelling the dose dependence of the effects of blend (Z)-3-hexenyl acetate and linalool on (A) response to pheromone and (B) pheromone salience. Circles = experimental values after normalization in linalool (black dots), (Z)-3 hexenyl acetate (green dots), and the 1:1 blend (blue dots). Lines = predicted values obtained from fits of the modified Hill’s equation 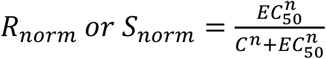 by a non-linear regression (see estimated parameter values in Table S2).

### Multi component blends reproduced the activity of salient compounds

Finally, we tested the effects of two more complex backgrounds on pheromone detection by Phe-ORNs. A 3-component background was prepared by mixing the agonist (Z)-3-hexenyl acetate with two inactive compounds, indole and β-caryophyllene, at a ratio of 1:1:0.3. A 4-component background was prepared by mixing the two agonists, (Z)-3-hexenyl acetate and linalool and the two inactive VPCs, α-pinene and eucalyptol, at a ratio of 1:1:2:2. The 2-component background obtained by mixing linalool and (Z)-3-hexenyl acetate was included as a reference. All blends and reference backgrounds were presented at (Z)-3-hexenyl acetate concentration = 1 AU. The 3-component background did not influence neuron firing and response to the pheromone any differently from (Z)-3-hexenyl acetate alone (pairwise PERMANOVA, P = 0.444, Figure 9B) or the 2-component blend (P = 0.908). Similarly, the 4-component background did not influence neuron firing and response to the pheromone any differently from either (Z)-3-hexenyl acetate alone (pairwise PERMANOVA, P = 0.0088, Figure 9B) or the blend of the two agonists (pairwise PERMANOVA, P = 0098, Figure 9). No significant difference was observed between the 3- and 4-component blends either (P= 0.310).

**Figure 9:**
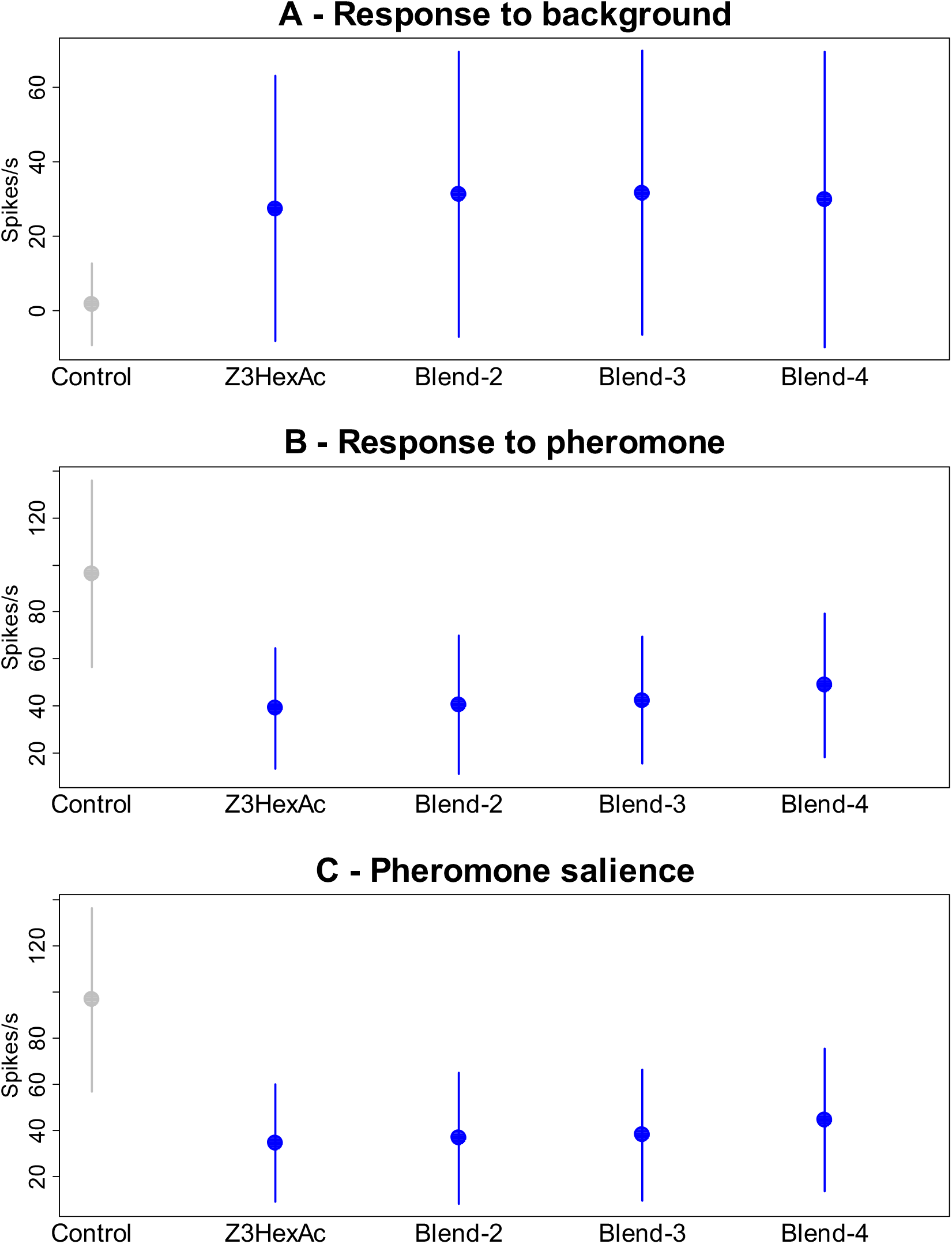
Effects of backgrounds with blends of 3- and 4-components on the Phe-ORN firing activity. Control = mineral oil. Z3HexAc = (Z)-3-hexenyl acetate. Blend-2 = (Z)-3-hexenyl acetate 1 AU plus linalool at ratios of 1:1. Blend 3 = (Z)-3-hexenyl acetate 1 AU plus indole and β-caryophyllene at ratios of 1:1:0.3. Blend 4 (Z)-3-hexenyl acetate 1 AU, linalool, α-pinene and eucalyptol at ratios of 1:1:2:2. (A) Response to the background (B) Response to the pheromone (C) Pheromone salience. Means of N = 34 measures on 17 Phe-ORNs. Error bars = standard deviations.

## Discussion

Our data confirm previous observations that a background of common VPCs interfere with the moth pheromone system in a dose-dependent manner. Interference with the neuronal coding of the pheromone signal starts at the periphery: specialized Phe-ORNs respond to some VPCs and their firing responses to the sex pheromone compound Z7-12:Ac are affected by a VPC background. Such interactions between odorants can potentially occur at many levels in the olfactory system, but the precise mechanisms have rarely been identified. In rats, non-competitive interactions resulting in mixture suppressions play a major role in the blend interactions that contribute to the perception of natural odorant mixtures (Rospars, Lansky et al. 2008). Competitive binding at the OR level has been firmly established in insects for the HR13 pheromone receptor of *Heliothis virescens* (Pregitzer, Schubert et al. 2012). In our electrophysiology experiments we mostly observed a reduced response to pheromone when it was presented over a single-VPC background. By contrast, in Ca-imaging, the response to the pheromone was slightly increased in a VPC background. This contrasts with studies in bees where Ca-imaging revealed a majority of mixture suppression effects in the AL (Deisig, Giurfa et al. 2006), but with a different protocol, COVs being delivered simultaneously to the bee antenna. In the present study while the firing response to VPCs decreased rapidly to a lower level after an initial peak, the calcium-response stayed at the same level for the whole duration of the background application to slowly decrease when it was turned off. This slower kinetics of the fluorescence measures do not reflect the fast change in firing frequency. To fully explain the decreased response to pheromone it should be noted that in our experiments the Phe-ORNs reached a certain degree of adaptation before the pheromone puff because of their response to the VPC background. Thus, in addition to a mixture suppression, cross-adaptation between background and pheromone probably also contributed to reduce the response to the pheromone. Whether environmental VPC concentrations can induce similar adaptation remains unknown.

VPCs differed in their capacity to stimulate Phe-ORNs and such differences cannot be explained by stimulus intensities because we adjusted the source concentrations to vapor pressures. Thus, these differences must be attributed to the binding selectivity of ORs or any other olfactory proteins. Although in the present work the pheromone salience was lower with VPCs activating the Phe-ORNs, other studies revealed that responses to pheromone can also be decreased by VPCs with no intrinsic activity (Party, Hanot et al. 2009). A clear inhibition of the pheromone response by linalool was observed *in vivo* in a noctuid moth, *Spodoptera littoralis*, although at the concentrations used linalool did not activate Phe-ORNs (Party, Hanot et al. 2009). This suggests that VPCs can alter the pheromone binding, whether it be by syntopic or allosteric interactions. It makes difficult to predict the impact of one VPC on the pheromone detection. Interestingly, we observed large differences between VPCs in the temporal dynamics of the firing responses. A fast response followed by a decrease was observed for (Z)-3-hexenyl acetate, while linalool elicited a delayed increase in firing. Differences in the time-course of odorant concentrations have been measured by physical methods in an olfactometer and they were attributed to air-surface interactions within the system delivering the stimuli (Martelli, Carlson et al. 2013, Gorur-Shandilya, Martelli et al. 2019). During transit odorants adsorb to the walls of the stimulator device and desorb later on, resulting in slower stimulus rise and decline, depending on the affinity of the odorant molecules for the tubing. It is not known whether a similar process also exists within olfactory organs and contributes to shape *in vivo* the stimulus course. In natural conditions, adsorption of volatiles like pheromones onto the vegetation is known to occur (Noldus, Potting et al. 1991). The physics of odorant transport adds a source of variability and a degree of complexity in the analysis of odor background effects.

Neuronal treatment of the input generally improves the signal-to-noise ratio in sensory-systems. The olfactory noise in natural environments is not only complex because of the diversity of VPCs but it fluctuates largely independently of the pheromone signal. Such an olfactory noise deteriorates intensity and quality coding of the pheromone signal in laboratory conditions (Renou, Party et al. 2015). Furthermore, because of their responses to VPCs, the Phe-ORN outputs may be ambiguous with respect to the nature of the chemical. Does the AL neuronal network facilitate signal identification in downstream olfactory areas by reformatting the ORN-input? The high convergence level of many ORNs expressing the same functional type of OR onto a few PNs averages the responses of many ORNs. However, we did not observe any increase in pheromone salience over background from Phe-ORNs to MGC-neurons. The ALs reformat the ORN output, resulting in increased signal-to-noise ratio (Bhandawat, Olsen et al. 2007, Masse, Turner et al. 2009). Convergence should increase pheromone signal to background ratio in PNs, provided not all Phe-ORNs have the same sensitivity to background odorants. However, if most Phe-ORNs respond in a standard way to a background odorant, averaging will not improve signal-to-noise ratio. Then, pooling Phe-ORN outputs might make more apparent among AL neurons the odor stimuli that produce weak ORN responses. To explain this contradiction, it is important to consider that separating signal from noise is not the only challenge of pheromone communication because pheromone concentrations vary over a very large range in natural conditions (Murlis and Jones 1981, Murlis 1986). The dynamic range of moth Phe-ORNs is of seven orders of magnitude while the maximum firing rate of individual neurons is in the range of 150 to 200 spikes per second (Rospars, Gremiaux et al. 2014). Neural gain control mechanisms (Root, Masuyama et al. 2008, Gorur-Shandilya, Demir et al. 2017) allow the brain to cope with large changes in the level of the ORN input. A number of studies suggests this is a key function of the AL (Wilson 2013). Gain control alters the relationships between ORNs and PNs firing in such a way that amplification is high when the ORN firing activity is weak, but low when it is strong. Such a non-linear amplification could also increase the representation in the MGC of weak agonists like VPCs. Thus, in the ALs, the neuronal network might have evolved to increase sensitivity and encode fast changes over a wide range of concentrations, possibly at some cost for qualitative selectivity.

VPCS are naturally emitted by plants in complex mixtures. While each VPC might individually be present at very low mixing rates within the atmosphere, the total levels of VPCs over a forest commonly reach hundreds of ppb to peak at nearly one ppm (Meneguzzo, Albanese et al. 2019). It is thus important to establish whether effects of VPC are cumulative, or whether VPCs interact with each other, and how much such interactions affect pheromone detection. To understand blend perception, interactions between odorants in mixtures have been intensively analyzed in various organisms, including insects, at the periphery (Munch, Schmeichel et al. 2013) and brain levels (Deisig, Giurfa et al. 2006, Shen, Tootoonian et al. 2013). In most cases the response to a blend is lower compared to the response to the most active component, an interaction called mixture suppression (Ache, Gleeson et al. 1988). However, none of these studies considered the case of a composite background interacting with detection of an odor signal by specialized receptors. Since the potential sites of interactions are multiple, interactions within multi-component blends are probably difficult to quantify. Nevertheless, dose-response relationships fitted quite well with classical Hill models, enabling quantitative comparison between blends and single compounds. Our analyses reveal that the complexity of the blend, in terms of its number of constituents, did not play a prominent role in the interaction with pheromone perception. Comparing three- or four-component blends to binary blends or single compound indicated that a blend showed the activity of its most active compound. Thus, although the diversity of a background might increase the probability of including a VPC capable of interacting with the pheromone system, chemical diversity of the background does not seem to be a prominent factor per se.

Insects have evolved their efficient pheromone communication system in the presence of a complex natural background of VPCs (Conchou, Lucas et al. 2019). However, among other anthropogenic factors, global warming is significantly affecting plant metabolism so that the emissions of VPCs are modified. Current knowledge on the impact of CO_2_ concentration and temperature elevations on plant physiology indicates a fast global increase in their VPC emissions (Holopainen, Virjamo et al. 2018) and significant changes in the pattern of terpenoid release (Ghirardo, Lindstein et al. 2020). This increase in atmospheric mixing rate of VPC will change olfactory landscapes which, as confirmed in our study, might impact pheromone communication. Also host plant location might be affected because it has been shown that the ratio between background and host-plant volatiles alter the response of female moths to their host-plant (Knudsen, Norli et al. 2017). Interestingly, differences between VPCs in their capacity to interfere with insect olfaction indicate that impact will greatly depend on which VPCs are concerned. Stimulation of the emission of indigenous plants, or introduction of novel plants that release salient VPCs, like for instance (Z)-3-hexenyl acetate for *A. ipsilon*, will have a greater impact on insect olfactory communication, than botanical changes associated to less potent VPCs. It is thus important to better evaluate the impacts of the exposure of insects to modified odorscapes. Analyses of interactions at the molecular level, will contribute to better predict the risks. We also need quantitative analyses of the composition of the odorscape at fine temporal and spatial scales to better estimate the VPC peak concentrations and exposure durations insects are exposed to.

## Acknowledgement

We acknowledge the technical assistance of Joanne Louison, Pascal Roskam, and Isabelle Touton, for insect rearing. We also thank Christelle Monsempès for sharing her expertise and training the first author in central extracellular electrophysiology.

