## supplemental figures for "Effects of an odor background on moth pheromone communication: constituent identity matters more than blend complexity"

### Conchou et al - Supplementary Information

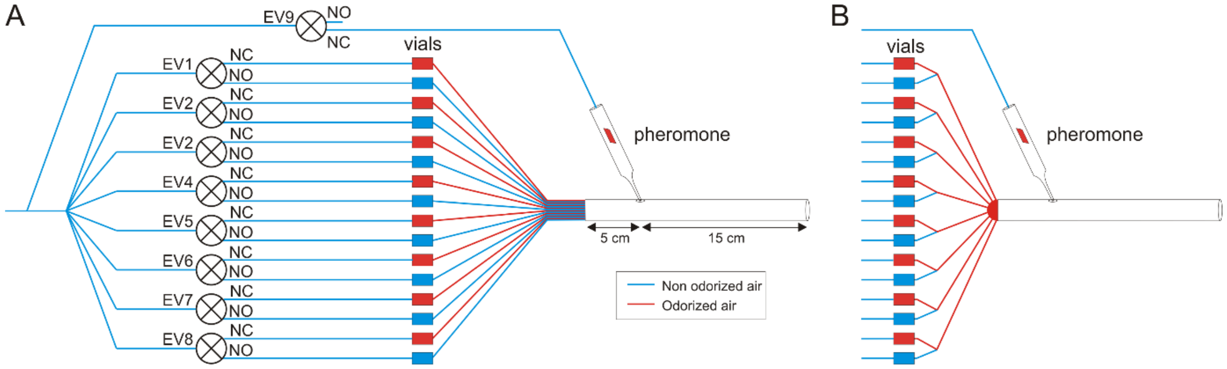


**Figure S1: Diagram of the stimulator devices.** **(A)** Complete diagram of the device used to create single VPC backgrounds. Vial sources contain only one VPC and each VPC lines are separated from each other up to the glass main tube where the pheromone is delivered. **(B)** Distal end of the stimulator used in the experiments using backgrounds with two or more VPCs. A low dead-volume manifold has been added to create a mixing chamber for VPCs before their entry in the glass tube. The upstream part (not represented) is identical to A. EV = electrovalve, NO = normally open exit, NC = normally closed exit. Line colors indicate when tubing conduct clean air (blue) and odorized air (red).

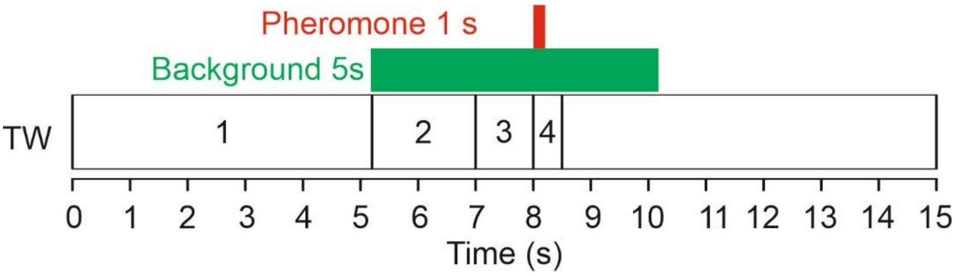


**Figure S2: Stimulation protocol for electrophysiological experiments on Phe-ORNs and MGC-neurons.** Green and red boxes indicate the delivery of the VPC background and pheromone on the moth antenna, respectively. TW1 to TW4: limits of the time windows used in the data analysis.

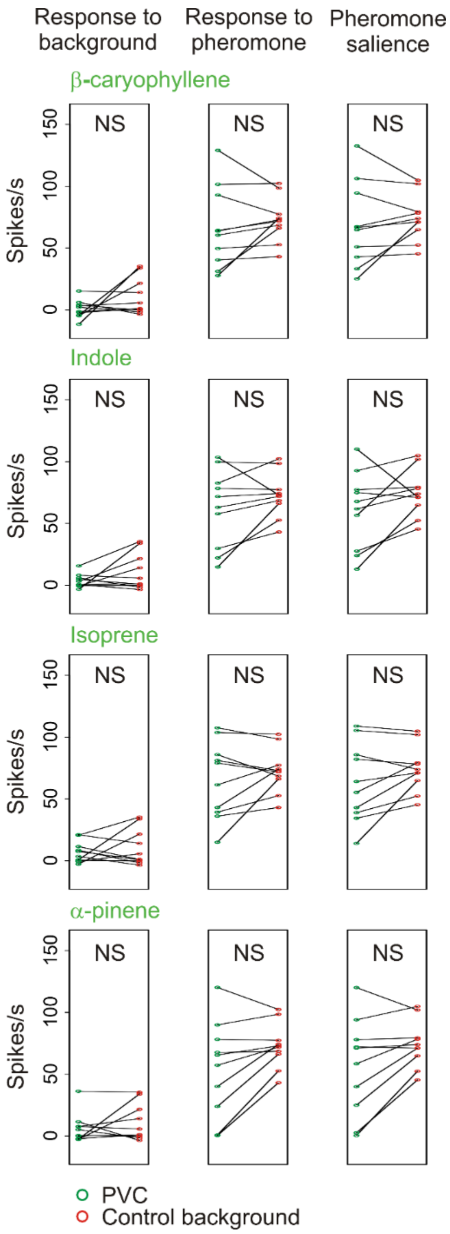


**Figure S3: Four VPCs, β-caryophyllene, indole, isoprene and α-pinene, neither activated MGC neurons nor modified their responses to the pheromone**. Strip charts compare individual neuron firing activities in each VPC background (green dots) with the control background (red dots). The firing frequency of MGC neurons was measured during different time windows to evaluate their response to background (left column), response to pheromone (middle column), and pheromone salience (right column). NS = p-value above FDR threshold. N = 10.

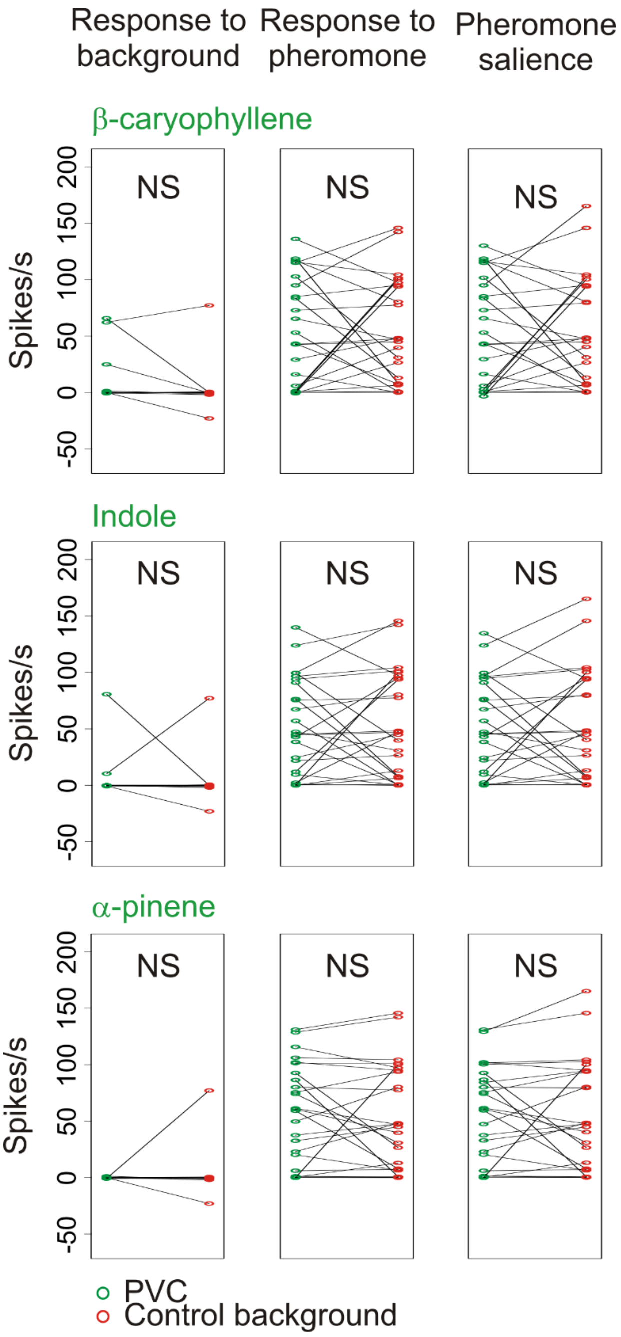


**Figure S4: Four VPCs, β-caryophyllene, indole, isoprene and α-pinene neither activated Phe-ORNs not modified their responses to the pheromone**. Strip charts compare individual neuron firing activities in each VPC background (green dots) with the control background (red dots). Phe-ORN firing frequency was measured during appropriate time windows to evaluate their response to background (left column), response to pheromone (middle column), and pheromone salience (right column). NS = p-value above FDR threshold. N = 26.

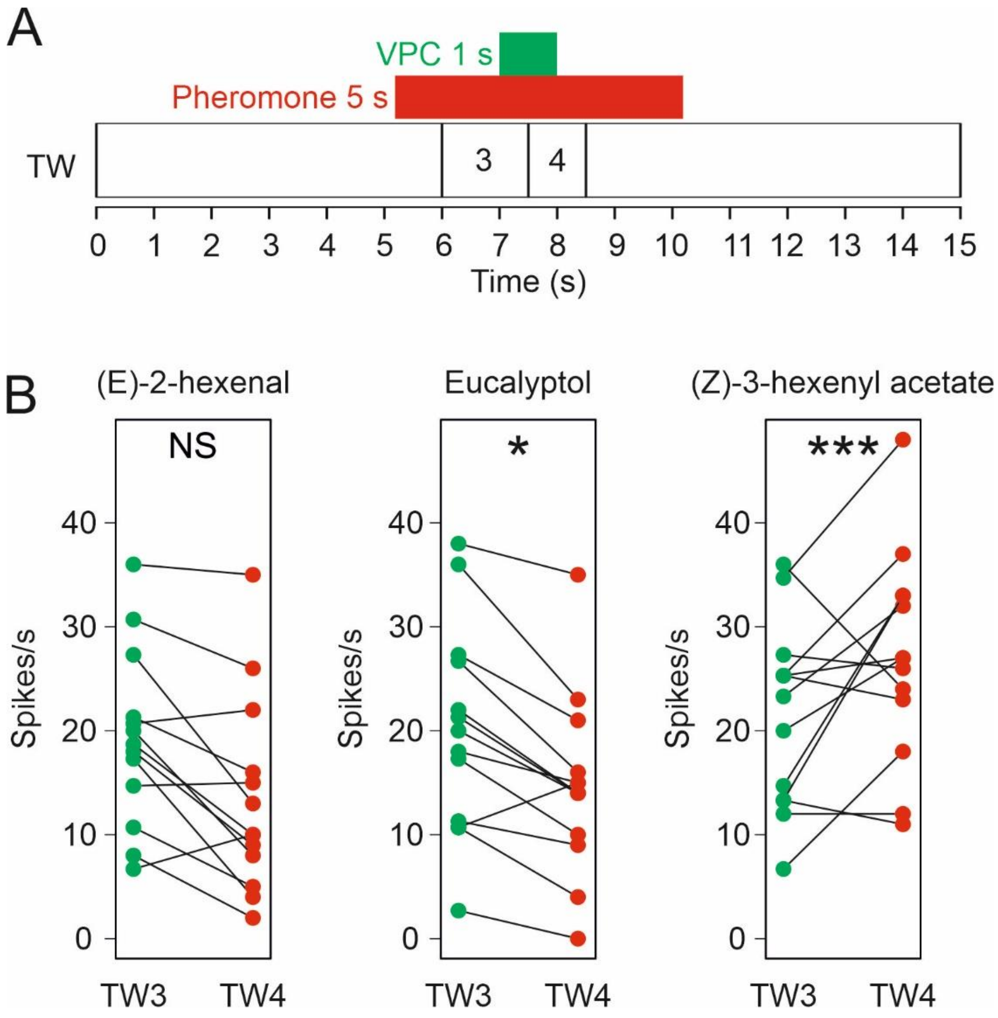


**Figure S5: A puff of eucalyptol has an inhibitory effect on the response to pheromone of Phe-ORNs.** (A) Stimulation protocol applied in this experiment. Green and red boxes indicate the delivery of the VPC background and the pheromone on the moth antenna, respectively. TW: limits of the two time windows used in the data analysis. (B) Comparison of the effects of a VPC puff (green dots) vs. control (red dots) on the pheromone response. N =13. Stars indicate p-values of the paired t test below FDR threshold.
