## supplemental tables 1 and 2 for "Effects of an odor background on moth pheromone communication: constituent identity matters more than blend complexity"

### Supplementary information: tables

| Volatile plant compound | CAS number | Nominal purity | Concentration in source (% v/v in mineral oil) | Partition coefficient $K_{hl}$ | Concentration in headspace (µmol/L for 1% v/v in mineral oil) |
|---|---|---|---|---|---|
| Indole | 120-72-9 | 99 % | saturated solution | $1.9 * 10^{-6}$ | 0.20 |
| (E)-2-hexenal | 6728-26-3 | 98 % | 0.1 % | $6.3 * 10^{-4}$ | 53.79 |
| Eucalyptol | 470-82-6 | > 98 % | 1 % | $1.4 * 10^{-5}$ | 0.84 |
| α-pinene | 80-56-8 | 98 % | 1 % | $3.5 * 10^{-5}$ | 2.20 |
| Linalool | 78-70-6 | 97 % | 1 % | $6.7 * 10^{-6}$ | 0.38 |
| Isoprene | 78-79-5 | > 99.5 % | 0.0001 % | nd | nd |
| β-caryophyllene | 87-44-5 | > 80 % | 10 % | $2.3 * 10^{-7}$ | 0.01 |
| (Z)-3-hexenyl acetate | 3681-71-8 | > 98% | 1 % | $2.6 * 10^{-5}$ | 1.64 |

**Table S1**: **List of the VPCs used to produce odorant backgrounds** with their air-mineral oil partition coefficients ($K_{hl}$) and estimated concentrations in source background.

| Response to pheromone | | | | | | | | |
|---|---|---|---|---|---|---|---|---|
| | EC | | | | n | | | |
| | | sd | t value | P | | sd | t value | P |
| (Z)-3-hexenyl acetate | 0.4654 | 0.1066 | 4.364 | $4.920*10^{-5}$ | 0.6585 | 0.1312 | 5.018 | $4.67*10^{-6}$ |
| linalool | 1.8279 | 0.7733 | 2.364 | $2.123*10^{-2}$ | 0.4458 | 0.1303 | 3.421 | $1.11*10^{-3}$ |
| blend | 0.6427 | 0.1041 | 6.174 | $5.650*10^{-8}$ | 0.7339 | 0.1056 | 6.950 | $2.62*10^{-9}$ |
| **Pheromone salience** | | | | | | | | |
| (Z)-3-hexenyl acetate | 0.4228 | 0.0806 | 4.913 | $6.870*10^{-6}$ | 0.7790 | 0.1472 | 5.293 | $1.67*10^{-6}$ |
| linalool | 1.5073 | 0.5678 | 2.654 | $1.008*10^{-2}$ | 0.4769 | 0.1323 | 3.606 | $6.22*10^{-4}$ |
| blend | 0.6001 | 0.0876 | 6.852 | $3.870*10^{-9}$ | 0.8519 | 0.1183 | 7.201 | $9.65*10^{-10}$ |

**Table S2: Estimation of the parameters of the modified Hill equations** used for modeling of dose-response relations for response to pheromone and pheromone salience in (Z)-3-hexenyl acetate, linalool or blend backgrounds. Pooled data (n = 16) were fitted to equations $R_{norm}\ or\ S_{norm} = \frac{EC_{50}^{n}}{C^{n}+EC_{50}^{n}}$ (see material and methods) by a non-linear regression.
